# *Bacillus velezensis* stimulates resident rhizosphere *Pseudomonas stutzeri* for plant health through metabolic interactions

**DOI:** 10.1101/2021.06.02.446779

**Authors:** Xinli Sun, Zhihui Xu, Jiyu Xie, Viktor H. Thomsen, Taimeng Tan, Mikael L. Strube, Anna Dragoš, Qirong Shen, Ruifu Zhang, Ákos T. Kovács

## Abstract

Trophic interactions play a central role in driving microbial community assembly and function. In gut or soil ecosystems, successful inoculants are always facilitated by efficient colonization, however, the metabolite exchanges between inoculants and resident bacteria are rarely studied, particularly in the rhizosphere. Here, we used bioinformatic, genetic, transcriptomic and metabonomic analyses to uncover syntrophic cooperation between inoculant (*Bacillus velezensis* SQR9) and plant-beneficial indigenous *Pseudomonas stutzeri* in the cucumber rhizosphere. We found that the synergistic interaction of these two species is highly environmental dependent, the emergence of syntrophic cooperation was only evident in a static nutrient-rich niche, such as pellicle biofilm in addition to the rhizosphere. Our results identified branched-chain amino acids (BCAAs) biosynthesis pathway involved in syntrophic cooperation when forming coculture biofilms. Assaying the metabolome further demonstrated metabolic facilitation among the bacterial strains. In addition, biofilm matrix components from *Bacillus* were essential for the interaction. Importantly, the two-species consortium promoted plant growth and helped plants alleviate salt stress. In summary, we propose a mechanism in which synergic interactions between a biocontrol bacterium and a partner species promote plant health.

## Introduction

Plants host enormous diverse communities of microorganisms, the plant microbiome, which is crucial for plant health. Beneficial plant-microbiome interactions improve plant fitness through growth promotion, stress alleviation, and defense against pathogens through various mechanisms [1]. Direct stimulations are mediated through production of phytohormones or 1-aminocyclopropane-1-carboxylate deaminases, induction of systematic resistance, and increasing nutrient acquisition through nitrogen fixation, phosphorus solubilization, and secretion of siderophores [2]. Indirect stimulation includes suppression of pathogens by antibiotic production, competition for niches within the rhizosphere, promotion of mycorrhizal functioning and changing the rhizosphere microbial community structure [3–5]. However, the high complexity of microbiome composition makes it challenging to answer the fundamental ecological questions surrounding natural microbial communities. Reductionist approaches conducted under laboratory conditions have been a promising strategy to decipher the microbial interactions in the rhizosphere microbiome along with their relevance for host health [6]. Driven by great application potential, studies focusing on synthetic communities (Syncom) have emerged in the last decade providing sustainable agriculture solutions [7]. Syncom are a microbial community designed by mixing selected strains to demostrate plant-beneficial impact. Such approach enables detailed assessment of host and microbe characteristics under controlled, reproducible conditions.

Two fundamental principles are commonly used for designing Syncom, function- and interaction-based approaches, during which cell-cell interactions play a key role in stability and robustness [7]. Given the high complexity of metabolites in the rhizosphere, metabolite exchange possibly drives species interactions in the plant microbiome. A well-studied example of positive microbial interactions includes cross-feeding, i.e., the exchange of essential nutritional molecules [8]. Gut microbiota studies highlighted that obligate cross-feeding can significantly expand the ecological niche space of each member involved in Syncom [9]. Moreover, cross-feeding also helps maintaining the diversity and stability of natural microbial communities [10], while spatial structure, especially biofilm, is vital for cross-feeding development and its evolution [11]. However, compared with the plentiful knowledge from gut microbiota studies, relatively few studies have been published on metabolite cross-feeding in the rhizosphere. We propose to exploit such metabolic mutualisms for designing robust probiotic Syncom for green agriculture.

Generally, *Bacillus* spp. and *Pseudomonas* spp. are the most extensively studied beneficial microorganisms in the rhizosphere [12, 13], several commercial products belong to these two genera are currently available for agricultural production of crops. Even though the biocontrol ability and plant-growth promotion by plant growth promoting rhizobacteria (PGPR) have been investigated thoroughly, their impacts on the composition of indigenous rhizosphere microbiome are still not fully explored. Microbial inoculants were shown to recruit assemblages of beneficial taxa, like *Flavobacterium*, *Pseudomonas, Agrobacterium*, and *Lysobacter* [14, 15], and inhibit soil-borne pathogens. It has been suggested that in addition to direct association with host plants, disease suppression and growth promotion may also be achieved by recruitment of beneficial species that reshape soil microbiome structure and function. However, these findings were largely constrained by microbiome sequencing approach, with limited information on the activity of the organisms behind them. We propose that the undiscovered interaction between PGPR and recruited beneficial microbes affect the functional capacities of the applied inoculants in the rhizosphere. Instead of focusing on the mono-association with plant, more attention should be paid to how inoculants modulate the structure of indigenous microbiome and how microbial interaction affect the functionality of the applied inoculants [16].

To address this important knowledge gap, we used *B. velezensis* SQR9 as a representative of PGPR to characterize the influence of PGPR on resident rhizosphere microbiome and study emerging microbial interactions using a simplified two-species community with target rhizosphere community bacteria. Plant beneficial *Bacillus*, including *B. velezensis* SQR9 are known for plant growth promotion, disease suppression and enhanced salt stress tolerance [17–20]. Root colonization and plant growth promoting properties require efficient biofilm formation on the roots. The essential *B. velezensis* genes to produce biofilm extracellular matrix include the *epsA-O* operon (encoding exopolysaccharide EPS) [21] and the *tapA-sipW-tasA* operon (encoding TasA protein fibers) [22]. Inoculation of corresponding bio-fertilizer to soil increased the abundance of indigenous microbial groups with reported antifungal activity, such as *Lysobacter* spp., which could play a keystone role in soil suppressiveness [14]. However, the exact mechanisms involved in the recruitment have not been characterized in detail previously. In this study, we discovered that *B. velezensis* SQR9 stimulated resident rhizosphere member, *P. stutzeri*. These two microorganisms formed robust biofilms *in vitro* and on the plant root surface. Metabolic and transcriptome analysis revealed potential cross-feeding to increase community performance. We demonstrated that the BCAAs biosynthesis pathway involved in syntrophic interaction of these two species during forming coculture biofilms. Furthermore, a Syncom containing these two strains excelled in plant growth promotion compared to the individual species. Together, our work expanded the knowledge on complex microbial interactions that generally occur in the rhizosphere, providing guidance for rhizosphere engineering in safe and eco-friendly agriculture.

## Methods and materials

### Rhizosphere sample collection

This study focused on cucumber rhizoshphere bacterial communities. The soil used in this study was collected from a field in Taizhou city, Jiangsu province, China (32.4555° N, 119.9229° E) in October 2017. Cucumber seeds (Jinchun 4) were purchased from Jiangsu Academy of Agricultural Sciences. The seeds were surface-sterilized and grown in ¼ Muarshige Skoog (MS) medium [23] for two weeks and then transferred to 2 kg soil pre-inoculated with 10 mL *B. velezensis* SQR9 suspensions (10^8^ cells mL^-1^). Plants were grown in a greenhouse at 30 °C, 16 h light /8 h dark. No inoculating soil was set as control. After 16 days, rhizosphere soil was collected as described by Bai [24]. Each treatment had 6 replicates.

### Strain isolations, culture conditions and mutant construction

Bacterial strains and plasmids used in this study are listed in Table S1. *B. velezensis* strain SQR9 (CGMCC accession number 5808) was isolated previously from cucumber rhizosphere, and used throughout this study. Deletion mutants were generated by using a markerless deletion method described by Zhou [25]. Oligonucleotides used for PCR in this study are listed in Table S2.

To isolate cooperating bacteria, fresh cucumber rhizosphere soil inoculated with strain SQR9 was collected and suspended in PBS buffer, and vortexed vigorously. Dilution series were plated on 0.1X TSB, R2A, TYG, and M715 media [24] solidified with 1.5% agar, then incubated at 30 °C for 2-7 days. 267 colonies were picked and phylogenetically characterized by colony PCR using a universal primer set (27F and 1492R) for the 16S rRNA gene and stored at −80 °C in 25% (v/v) glycerol.

*P. sturzeri* XL272 was selected in the following investigation and modified with a mini-Tn7 transposon containing a dsRed marker [26]. The genome was sequenced using a PacBio RS II platform and Illumina HiSeq 4000 platform at the Beijing Genomics Institute (BGI, Shenzhen, China). Sequences of the genome were deposited at the National Center for Biotechnology Information (NCBI) under Nucleotide accession number NZ_CP046538.

### Microbiome analysis

Total genomic DNA of soil samples was isolated using the PowerSoil® DNA Isolation Kit (Mo Bio Laboratories, Inc., Carlsbad, CA, USA). Mixed universal primers targeting the V3-V4 regions of 16S rRNA gene were used to construct the DNA library for sequencing. Paired-end sequencing of bacterial amplicons were performed on the Illumina MiSeq instrument (300 bp paired-end reads). Raw sequencing data have been deposited to the NCBI Sequence Read Archive (SRA) database under BioProject accession number PRJNA727458. Reads were processed using the UPARSE pipeline [27]. The paired-end reads were merged using the “fastq_mergepairs” command. High-quality sequences were then selected using the “fastq_filter” command and dereplicated using the “derep_fulllength” command. The singletons were removed using USEARCH-unoise3 algorithm and chimeric sequences were removed using “uchime_ref” command. The remaining sequences were used to create ASV table. Taxonomy assignment was performed using the Ribosomal Database Project (RDP) classifier. Bray-Curtis distance-based PCoA analysis, and permutational multivariate analysis of variance (PERMANOVA) were performed based on the ASV table using the vegan R package. Welch’s t test was used to calculate the significance of differences between two treatments using STAMP [28]. ASVs whose relative abundance were higher than 0.05% in all samples were used in this analysis.

### Biofilm formation assay

To observe the cocultured colony biofilm, equal volume of *B. velezensis* and *P. stutzeri* were mixed at an OD_600_ of 0.0001, then 5 μL of bacteria were spotted on TSB medium solidified with 1.5% agar. The plates were incubated at 30 °C for 48 h.

To observe pellicle biofilm formation, 20 μL of the start inoculum was cultivated in 2 mL of TSB liquid medium in a 24-well microtiter plate (Fisher Scientific). The microtiter plates were incubated statically at 30 °C for 24 h. The start inoculum was obtained by growing the cells overnight to exponential growth in TSB medium at 30 °C, 180 rpm shaken condition, the cells were spined down and diluted to OD_600_ of 1 in 0.9% NaCl buffer. For coculture, the start inoculum was prepared by mixing equal volume of two species. Without specific statement, the start inoculum was prepared as described here in all the experiments.

### Biofilm biomass quantification assay

Pellicle was grown in 6-well microtiter plates insert with 100 um Sterile Nylon Mesh Cell Strainers (Biologix Cat#15-1100). 10 mL of TSB liquid medium and 100 μL of the start inoculum were added. The plates were incubated for 24 h at 30 °C statically to allow the pellicle to grow on top of the nylon mesh cell strainer. The cell strainer was taken out, removed visible drops with paper, and weighed. The fresh weight was the total weight minus the weight of the nylon mesh. The dry weight was measured by drying the pellicle within the laminar hood for 24 h. Each treatment had 6 replicates.

### Cell numbers quantification

Cell numbers in coculture were quantified under four conditions: static TSB medium, shaken TSB medium, static MSgg medium [29], static root exudate medium (REM). Root exudates were collected as described by Feng [30]. REM was composed of 0.02 mg/mL root exudates and M9 glucose medium. M9 glucose medium was prepared from M9 minimal salts (5x, Sigma-Aldrich M6030), 2 mM MgSO_4_, 0.1 mM CaCl_2_ and 10 mg/mL glucose. For quantifying individual cell numbers in pellicle, 100 μL of the start inoculum were grown in 6-well microtiter plates (VWR) with 10 mL medium. A 100 μm sterile nylon mesh cell strainer and a Spectra Mesh™ Woven Filter (Fisher Scientific, Spectrum™ 146488) were put inside. The mesh was manually cut into 1.5 cm^2^ squares and autoclaved. After 48 h of pellicle development, the nylon mesh cell strainer was taken out, the inner filter was transferred to a 1.5 mL microcentrifuge tube, stored at −80 °C for following DNA or RNA extraction. This ensured equal sampling of biofilm. For quantifying individual cell numbers in shaken TSB, 40 μL of the start inoculum were inoculated in sterile test tubes with 4 mL TSB medium and incubated at 30 °C, 180 rpm shaken condition. After 48 h, 2 mL of cell cultures were spined down and stored −80 °C. Total DNA of biofilm formed on the mesh filter or cell pellets was extracted with E.Z.N.A.® Bacterial DNA Kit (Omega Bio-tek, Inc.) according to the manufacturer’s instructions.

Alignment of the *B. velezensis* SQR9 and *P. stutzeri* XL272 genomes was conducted with Roary [31] to find different genes of the two isolates. The strain-specific primer pairs for qPCR were designed based on different gene sequences according to the guidelines of Oligo (v7). The specificity of obtained primers was checked by conventional PCRs and qPCR melt curves. Standard curves were generated using plasmids containing corresponding fragments. qPCR was performed with Applied Biosystems (ABI) Real-Time PCR Instrument. Reaction components are as follow: 7.2 μL H_2_O, 10 μL 2× ChamQ SYBR qPCR Master Mix (Vazyme), 0.4 μL 10 μM of each primer and 2 μL template DNA. The PCR programs were carried out under the following conditions: 95 °C for 10 min, 40 cycles of 95 °C for 30 s, 61 °C for 30 s and 72 °C for 40 s, followed by a standard melting curve segment. Each treatment had 6 replicates, and each sample was run in triplicates.

### Whole-genome transcriptomic analysis and qRT-PCR validation

Total RNA was obtained from biofilms formed on Spectra Mesh™ Woven Filter in static TSB medium using the E.Z.N.A. bacterial RNA kit (Omega Bio-tek, Inc.), according to the instructions. Pair-end reads libraries were generated using NEBNext® Ultra™ Directional RNA Library Prep Kit for Illumina® (NEB, USA) and sequenced on an Illumina Hiseq platform. Raw sequencing data have been deposited to the NCBI SRA database under BioProject accession number PRJNA727814. Raw reads were quality-trimmed and then mapped to reference genomes using Bowtie 2-2.2.3 software [32]. Differential expression analysis was performed using the DESeq2 R package [33]. The resulting p-values of genes were adjusted using Benjamini and Hochberg’s approach for controlling the false discovery rate (FDR). Genes were assigned as differentially expressed when log_2_ fold change (LFC) >2 and FDR <0.05. For functional analysis, the protein-coding sequences were mapped with KEGG Orthology terms using EggNOG-mapper v2 [34]. P-values of pathways were corrected for multiple hypothesis testing using the Benjamini and Hochberg’s approach.

Isolated RNAs were reverse transcribed into single-stranded complementary DNA (cDNA) using the PrimeScript RT reagent kit with a genomic DNA (gDNA) eraser (Toyobo). Transcript levels of *ilvA*, *ilvC*, *ilvD*, *ilvE*, *ilvH*, *leuA*, *leuB*, *leuC* and *leuD* were measured by qRT-PCR using a SYBR Premix Ex Taq (perfect real time) kit (TaKaRa, Dalian, China). The *recA* gene was used as an internal control for *B. velezensis* SQR9. The *rpoD* gene was used as an internal control for *P. stutzeri* XL272. ABI Real-Time PCR Instrument was operated under the following conditions: cDNA was denatured for 10 s at 95 °C, followed by 40 cycles consisting of 5 s at 95 °C and 34 s at 60 °C. The relative expression of specific genes was calculated by using the 2−^ΔΔCT^ method [35].

### Growth curve assay

*B. velezensis* SQR9 and *P. stutzeri* XL272 were grown in TSB rich medium individually for 24 h, the cell cultures were spined down, then the supernatants were filter sterilized as bacterial metabolites. 2 μL of start inoculums were inoculated to 200 μL TSB medium or TSB supplemented with 10% bacterial metabolites in a 10×10 well Honeycomb Microplate. OD_600_ was measured every 30 minutes at 30 °C with Bioscreen C Automated Microbiology Growth Curve Analysis System.

### Metabolic facilitation assay and metabolome analysis

Both isolates were inoculated individually in 100 mL of M9 glucose medium, then incubated at 30°C, 180 rpm shaken condition. The consumption of glucose was measured every day using the Glucose GO Assay Kit (Sigma). As a result, *B. velezensis* SQR9 consumed all the glucose provided in 6 days, while *P. stutzeri* XL272 took 4 days. The cell cultures were spined down, then the spent medium was filter sterilized. Each isolate was inoculated in 20 mL of each other’s spent medium at 1% v/v, then incubated at 30 °C, 180 rpm shaken condition for extra 4 days. Each treatment had 4 replicates.

Two rounds of extracellular metabolites were collected and analyzed by UHPLC-MS/MS. Untargeted metabolomics analysis was performed using a Vanquish UHPLC system (Thermo Fisher) coupled with an Orbitrap Q Exactive HF-X mass spectrometer (Thermo Fisher). The raw data files generated by UHPLC-MS/MS were processed using the Compound Discoverer 3.0 (CD 3.0, Thermo Fisher) to perform peak alignment, peak picking, and quantitation for each metabolite. After that, peak intensities were normalized to the total spectral intensity. The normalized data were used to predict the molecular formula based on additive ions, molecular ion peaks, and fragment ions. And then peaks were matched with the mzCloud (https://www.mzcloud.org/) and ChemSpider (http://www.chemspider.com/) database to obtain the relative quantitative results. The raw data were processed on the free online platform of Majorbio Cloud Platform (www.majorbio.com).

### Root colonization assay

Colonization of *Arabidopsis thaliana* roots was performed according to the protocol from Dragoš [36]. To access root biofilm productivities, the roots were transferred into Eppendorf tubes, subjected to standard sonication protocol and the CFU assays were performed for obtained cell suspensions. To extract CFU/mm of the root, the obtain CFU values were divided by the total length of a corresponding root.

### Greenhouse experimental design

The trial was conducted from June to August 2020, in the greenhouse of Nanjing Agricultural University. The soil used for pot experiments were collected from field sitewith a histroy of cucumber cultivation. The field site was located in Nanjing, Jiangsu Province, China and the soil had following properties: pH 5.62, organic matter 21.6 mg/kg, available N 157 mg/kg, available P 128 mg/kg, available K 268 mg/kg, total N 1.85 g/kg, total P 1.86 g/kg, total K 15.3 g/kg. Pots with 2 kg soil were divided into two group. Group A: untreated soil. Group B: salt-treated soil group, the required amount of NaCl was added into soil and stirred well to blend to attain to 3.00 g/kg salt concentration. After one week soil incubation, one-week-old cucumber seedlings were transfer into the soil and growth for another 5 days. Then, for each group, the experiment includes four treatments: CK, un-inoculated treatment; S, plants treated with 10 mL *B. velezensis* SQR9 suspensions (10^8^ cells/mL); P, plants treated with 10 mL *P. stutzeri* XL272 suspensions (10^8^ cells/mL); PS, plants treated with 5 mL *B. velezensis* SQR9 and 5mL *P. stutzeri* XL272 suspensions, respectively (10^8^ cells/mL). Plants were grown for another 30 days at 30 °C, 16 h light / 8 h dark. No inoculating soil was set as control. Each treatment had ten to twelve replicates.

### Plant growth promoting (PGP) traits detection

*P. stutzeri* XL272 was tested for PGP traits including production of indoleacetic acid (IAA), ammonia, siderophore and phosphate solubilization. IAA production and ammonia production were detected by the method as described in [37]. Siderophore production was detected on CAS agar [38]. Phosphate solubilization was detected on NBRIP agar contained calcium phytate or Ca_3_(PO_4_)_2_ [39].

### Microscopy/confocal laser scanning microscopy

Fluorescent images of colonies and whole pellicle were obtained with an Axio Zoom V16 stereomicroscope (Carl Zeiss, Jena, Germany) equipped with a Zeiss CL 9000 LED light source and an AxioCam MRm monochrome camera (Carl Zeiss) and HE eGFP (excitation at 470/40 nm and emission at 525/50 nm), and HE mRFP (excitation at 572/25 nm and emission at 629/62 nm) filter sets. The exposure times for green and red fluorescence were set up to maximal possible values before reaching overexposure, using the range indicator function.

The pellicles and root colonization were visualized with a confocal laser scanning microscopy (LMI-005-Leica Microsystems Confocal Microscope-SP8). Fluorescent reporter excitation was performed with the argon laser at 488 nm and 556 nm, the emitted fluorescence was recorded at 484–536 nm and 560-612 nm for GFP and DsRed, respectively. ImageJ software was used to obtain overlaid, artificially colored images for both stereomicroscope and CLSM.

### Statistical analysis

Analysis was conducted in R 4.0.3 and figures were produced using the package ggplot2 or GraphPad Prism 8. Detailed statistical analysis were described in the figure legends.

## Results

### *B. velezensis* SQR9 induce the enrichment of *Pseudomonas* spp. in the cucumber rhizosphere

To explore the effects of *B. velezensis* SQR9 on rhizosphere microbiota, two-weeks-old cucumber seedlings were inoculated with strain SQR9 and the rhizosphere soil samples were collected after sixteen days. 16S rRNA amplicon sequencing was applied to compare the composition of rhizo-microbiomes of untreated control plant and plant treated with *B. velezensis* SQR9. Principal coordinates analysis (PCoA) was used to visualize differences in taxonomic abundance using bray-curtis distances (Fig 1A). Untreated rhizosphere samples (CK) were clearly separated from the *B. velezensis* SQR9 inoculated samples (S), demonstrating that strain SQR9 had an influence on the indigenous bacterial community. To illustrate changes in the community composition and reveal the affected species at genus level, STAMP analysis was applied [28]. In total, 21 Amplicon sequence variants (ASVs) were significantly influenced by strain SQR9 (t test; p <0.05) (Fig 1B). Based on the differences in relative abundance, members of the genera *Pseudomonas*, *Vogesella*, *Pseudoxanthomonas*, *Chryseobacterium*, *Pseudoduganella*, *Lysobacter*, *Klebsiella* and *Cellvibrio* were increased after SQR9 application. Remarkably, eight out the twenty-one ASVs mapped to the *Pseudomonas* genus, suggesting a positive interaction with strain SQR9. Recent studies previously demonstrated that these two beneficial genera have the potential of interacting positively to enhance plant growth [40] and rescue their host plant from a sudden-wilt disease [41, 42]. We hypothesized that *B. velezensis* SQR9 may recruit and then synergistically interact with specific beneficial *Pseudomonas* spp. that contribute to plant growth promotion.

**Figure 1.**
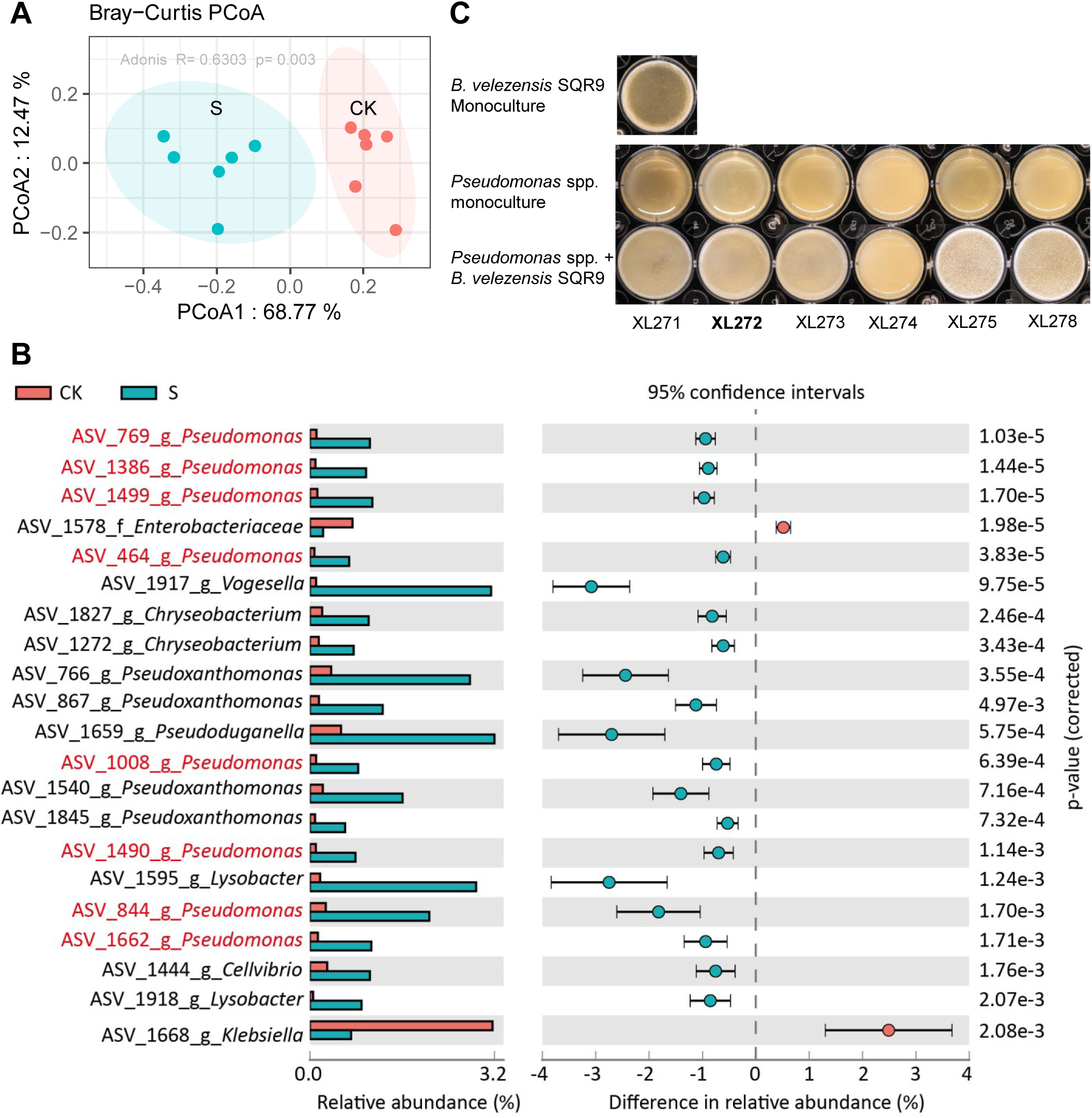
Influence of *B. velezensis* SQR9 on rhizosphere microbiota and biofilm phenotype of isolated bacteria with predicted synergism. **(A)** Principal Coordinates Analysis (PCoA) of the rhizosphere bacteria community are plotted based on the Bray-Curtis distance metrices for taxonomical data (p < 0.01). Permutational multivariate analysis of variance (PERMANOVA) were performed using the vegan R package. Samples were isolated from rhizosphere of untreated control (CK) and *B. velezensis* SQR9 inoculated (S) plants (n=6). **(B)** Differences in abundance of genera between control (CK) and *B. velezensis* SQR9 inoculated (S) rhizosphere samples (Welch’s t-test; p < 0.05; p value was corrected by Benjamini-Hochberg method). ASVs matched to *Pseudomonas* spp. are marked in red. **(C)** Biofilm phenotype of predicted cooperating strains. XL271-278 represent different *Pseudomonas* isolates. Well diameter is 15.6 mm.

To further characterize the potentially synergistic interaction between SQR9 and the recruited *Pseudomonas* spp., we isolated candidate bacteria from the rhizosphere of the *B. velezensis* SQR9 inoculated plants. In total, 267 bacterial isolates were obtained and phylogenetically characterized based on distinct 16S rRNA gene sequences (Fig S1), allowing us to recover six *Pseudomonas* isolates, for further interaction studies. Plant beneficial bacterial consortia have been suggested to form biofilm synergistically [41, 43]. Whether *Pseudomonas spp.* isolates act synergistically with *B. velezensis* SQR9 *in vitro* biofilm formation were tested. Intriguingly, five out of six *Pseudomonas* isolates showed enhanced biofilm phenotype in coculture as indicated by floating biofilm, pellicle surface complexity and biomass (Fig 1C & S2). The phenotype was especially pronounced in *P. stutzeri* XL272 where the coculture biofilm dry weight increased more than 3-fold (t test; p <0.01) compared to the SQR9 monoculture pellicle. This suggested that the recruited *Pseudomonas* species have positive interactions with *B. velezensis* SQR9.

### *P. stutzeri* XL272 form biofilm with *B. velezensis* SQR9 synergistically in rich medium but not in minimal medium

As *P. stutzeri* XL272 showed the highest synergy with *B. velezensis* SQR9 in biofilm formation, as indicated by enhanced biomass production when compared with that of single-species biofilm (Fig 2A, Fig S2), this isolate was selected for further experiments. This isolate has 98.06% sequence similarity to with ASV_1490, which is one of the increased ASVs. To illustrate whether the enhanced coculture biofilm resulted from cooperation or competition, the absolute cell numbers of the two species were quantified using qPCR. The cell numbers of both interaction partners were significantly higher in the dual-species biofilm (t test, p <0.01) in comparison with single-species biofilms in static TSB or REM, indicative of interspecies cooperation (Fig 2B& Fig S3A&D). The distribution of these two strains were also visualized under confocal laser scanning microscope using GFP tagged *B. velezensis* SQR9 and DsRed tagged *P. stutzeri* XL272. The two strains formed distinct cell clusters and segregated within the biofilm. *B. velezensis* SQR9 appeared as the dominant species in the population (Fig 2C), in line with qPCR quantification data (Fig. S3A). In a cocultured colony, the two strains also occupied different niches, with *P. stutzeri* at the bottom and *B. velezensis* on the top (Fig 2D). Altogether these results indicate that *B. velezensis* and *P. stutzeri* cooperate in biofilm mode potentially by niche partitioning.

**Figure 2.**
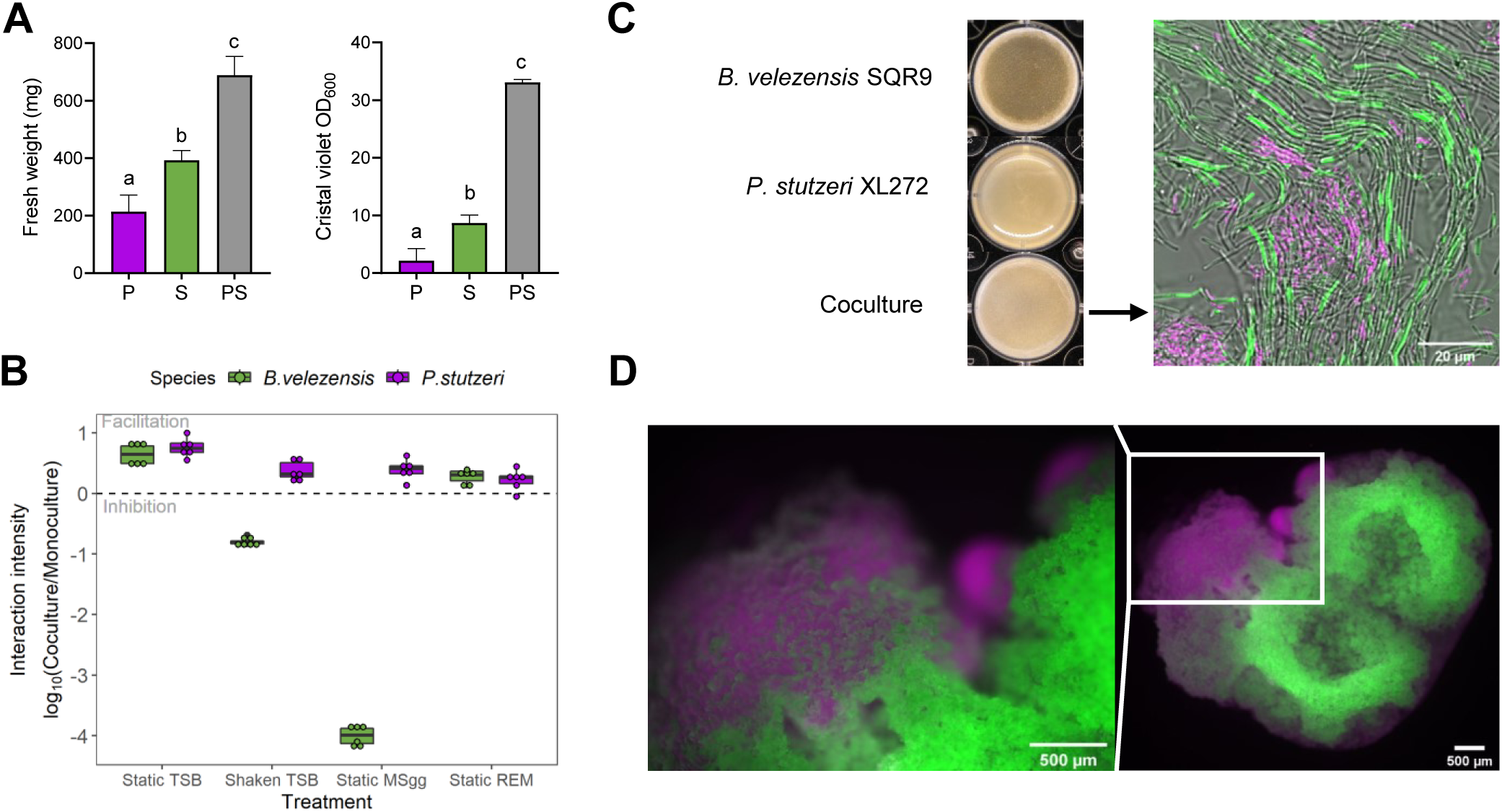
*P. stutzeri* XL272 and *B. velezensis* SQR9 form biofilm synergistically. **(A)** Biofilm biomass quantification by fresh weight and crystal violet assay. P represents *P. stutzeri* XL272 monocultured pellicle, S represents *B. velezensis* SQR9 monocultured pellicle, PS represents cocultured pellicle. Biofilm was cultivated in static TSB medium. Bars represent the mean ± s.d. (n = 6). Different letters indicate statistically significant (p < 0.05) differences according to ANOVA, Tukey test via Prism 8. **(B)** Interaction intensity is defined as logarithmic scale of DNA copies in cocultures relative to the average DNA copies in monocultures. Interaction intensity > 1 indicates facilitation, while < 1 indicates inhibition. Bars represent the mean ± s.d. (n = 6). Static TSB and static REM (root-exudate medium) represented structured, nutrient rich condition, shaken TSB represented unstructured, nutrient rich condition, static MSgg represented structured, nutrient limited condition. Cooperation only occurred under structured, nutrient rich condition. **(C)** Biofilm formation of monoculture and coculture. Biofilm formed by *P. stutzeri* XL272 (magenta) and *B. velezensis* SQR9 (green) were viewed under the CLSM. Well diameter is 15.6 mm. Scale bar represents 20 µm. **(D)** Colony grown on TSB agar. DsRed tagged *P. stutzeri* XL272 were colored in magenta, GFP tagged *B. velezensis* SQR9 were colored in green. Scale bar represents 500 µm.

Next, we tested the role of structured environment and nutrient availability of the cooperative relationship between *B. velezensis* SQR9 and *P. stutzeri* XL272. The interaction intensity was calculated as a logarithmic value of cell numbers in coculture relative to monoculture, where value > 1 indicates facilitation, while value < 1 indicates inhibition (Fig 2B). In the absence of spatial structure or under limited nutrient availability, only *P. stutzeri* was benefited in the coculture (Fig 2B& Fig S3B&C). While in the static, nutrient rich medium (TSB or REM), both species were facilitated by each other (Fig 2B& Fig S3A&D). These observations suggested that the mutualism between *B. velezensis* and *P. stutzeri* is only maintained under static, nutrient rich condition, further supporting our hypothesis on cooperation via niche partitioning.

### *B. velezensis* biofilm matrix EPS and TasA are required for the mutualism

To understand the molecular mechanisms of cooperation between *B. velezensis* and *P. stutzeri*, we examined the contribution of *B. velezensis* biofilm matrix components, EPS and TasA, for the interaction. In TSB medium, *B. velezensis* Δ*tasA* mutant showed severely impaired monocultured pellicle, but not Δ*epsD* (Fig 3A). To test whether these components contribute to the coculture pellicle phenotype, we mixed *P. stutzeri* strains with *B. velezensis* strains lacking either EPS or TasA. The resulting coculture pellicles appeared weaker than those formed by wildtype cocultures, wrinkled biofilms were not formed (Fig 3A). The cell numbers of both *B. velezensis* matrix mutants, were reduced in cocultures, indicating that EPS and TasA were both necessary for efficient mutualism (Fig 3B). Meanwhile, the cell numbers of *P. stutzeri* were unaffected in the different cocultures. We also tested interaction of *P. stutzeri* with *B. velezensis* matrix mutants in biofilm-inducing minimal medium MSgg. Surprisingly, although both *B. velezensis* mutants showed impaired individual fitness in minimal medium, they benefit from the presence of *P. stutzeri* (Fig S4). In conclusion, biofilm matrix components EPS and TasA are necessary for positive interspecies interaction.

**Figure 3.**
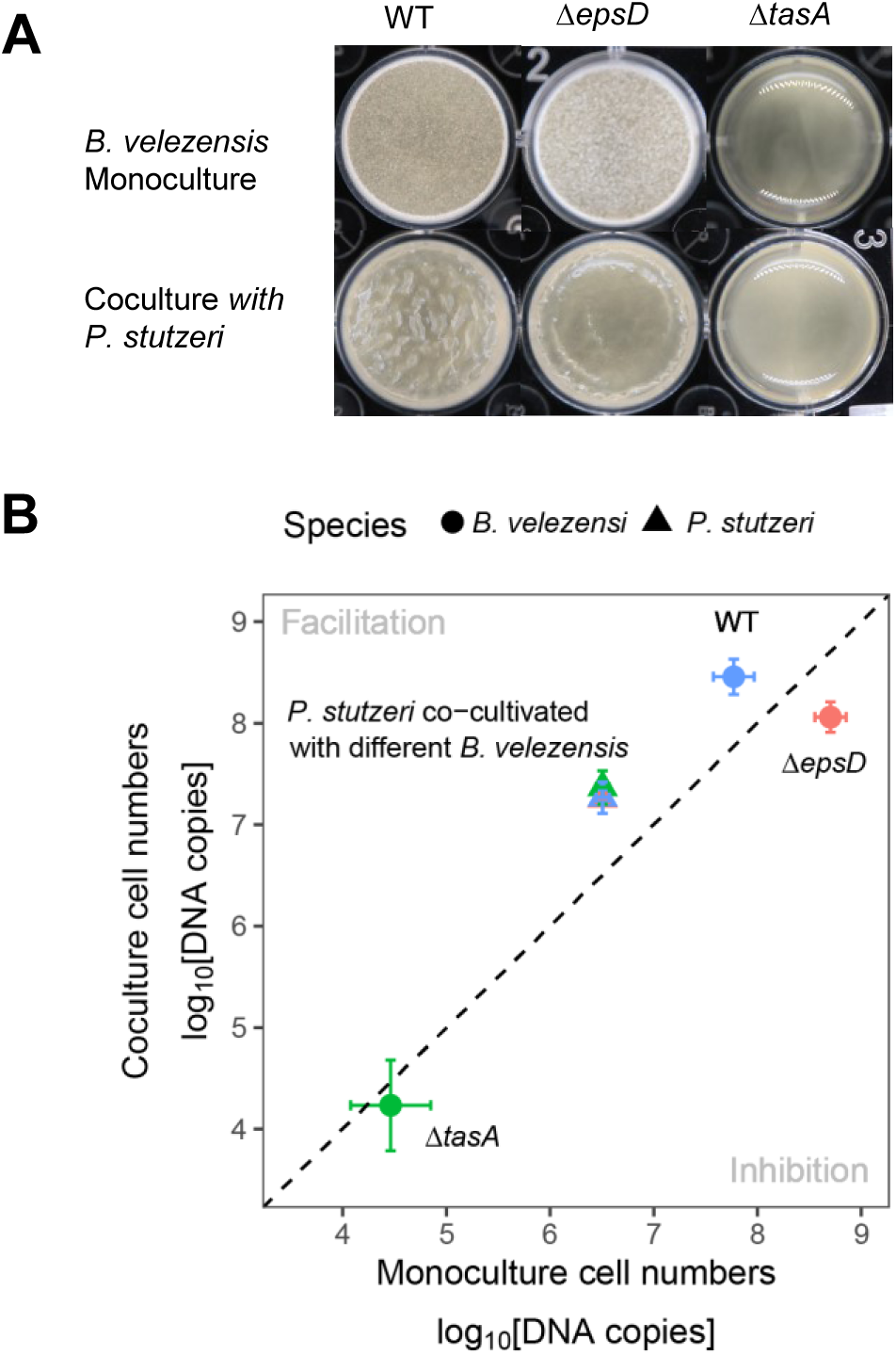
Extracellular matrix EPS and TasA are essential for mutualism in TSB rich medium. **(A)** Formation of pellicle biofilms by the mutants deficient in biosynthesis of exopolysaccharide EPS (Δ*epsD*) and TasA protein fibers (Δ*tasA*). Cells were incubated in TSB at 30 °C for 24h before images were taken. Well diameter is 15.6 mm. **(B)** Cell numbers in dual species biofilm. Circle dots represent *B. velezensis*, triangles represent *P. stutzeri*. Colors indicate *B. velezensis* in coculture or corresponding co-cultivated *B. velezensis,* WT (blue), Δ*epsD* (pink), Δ*tasA* (green). Data presented are the mean ± s.d. (n = 6). Error bars represent standard deviations.

### *P. stutzeri* XL272 might provide BCAAs to *B. velezensis* SQR9

Our results above demonstrated that *P. stutzeri* XL272 cooperate with *B. velezensis* SQR9 in TSB medium during biofilm formation. To disentangle the mechanism of cooperation, the transcriptomes of both species were determined in biofilm to elucidate the potential mode of actions underpinning the remarkably enhanced biomass in dual-species biofilm. In total, 345 genes of *B. velezensis* SQR9 and 443 genes of *P. stutzeri* XL272 were significantly regulated at a minimum of fourfold expression change in dual-species biofilms compared to single-species biofilms (24 h of interaction). Major transcriptional alterations were observed in genes related to bacterial metabolism, biosynthesis of amino acids, flagellar assembly, and ABC transporter (Fig. 4A). In the cocultures, fifteen *B. velezensis* SQR9 pathways were downregulated including amino acid biosynthesis and metabolism, sulfur metabolism, selenocompound metabolism, carbohydrate biosynthesis and metabolism, biosynthesis of secondary metabolites and ABC transporter (Fig. 4A). Intriguingly, the BCAAs biosynthesis pathway was downregulated in *B. velezensis* SQR9. qRT-PCR assay validated that all the genes involved in this pathway were downregulated in *B. velezensis* SQR9 while conversely upregulated in *P. stutzeri* XL272 during co-culturing (Fig 4B). This result suggested that the reduced production of BCAAs in *B. velezensis* SQR9 might be compensated by *P. stutzeri* XL272.

**Figure 4.**
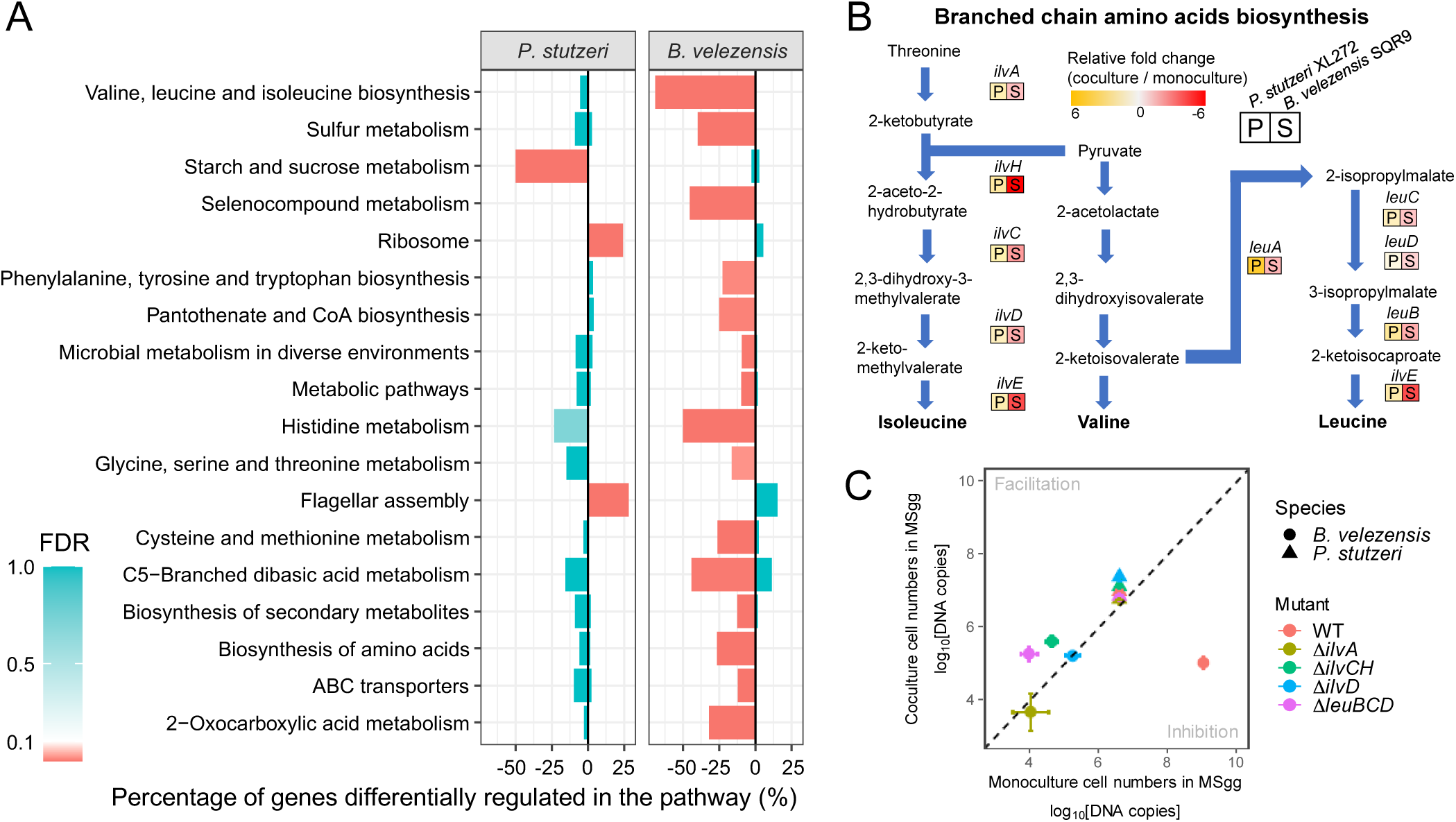
Transcriptional response of *P. stutzeri* XL272 and *B. velezensis* SQR9 in dual-species biofilm. A). KEGG pathway analysis of the genes induced (right bars) and repressed (left bars) in response to co-culturing. P represents *P. stutzeri* XL272, S represents *B. velezensis* SQR9. Rich factor indicated proportion of differentially expressed genes (LFC > 2, FDR < 0.05) in specific pathway. The resulting P-values were corrected for multiple hypothesis testing. B). The relative fold change of genes involved in the branched-chain amino acid biosynthesis pathways examined by qRT-PCR. Color represents the mean relative fold change of mRNA in coculture compared to monoculture (n = 6). C). Cell numbers of dual species biofilm in MSgg medium. Circle dots represent *B. velezensis*, triangles represent *P. stutzeri*. Colors indicate *B. velezensis* in coculture or corresponding co-cultivated *B. velezensis* WT (pink), Δ*ilvA* (brown), Δ*ilvCH* (green), Δ*ilvD* (blue), Δ*leuBCD* (magenta). *P. stutzeri* XL272 promote the growth of *B. velezensis* SQR9 Δ*ilvD* and Δ*leuBCD* in nutrient limited condition. Data presented are the mean ± s.d. (n = 6). Error bars represent standard deviations.

We further constructed *B. velezensis* BCAA biosynthetic mutants: Δ*ilvA*, Δ*ilvCH*, Δ*ilvD* that were unable to synthesize all three BCAAs, and Δ*leuBCD* that was unable to synthesize leucine. All the mutants showed severe growth defects in MSgg minimal medium under both planktonic and static conditions (Fig S5A-C). However, these strains showed increased static biofilm formation and higher planktonic biomass than wildtype in TSB, indicating that BCAAs can be provided exogenously (in rich TSB medium) (Fig S5A-C). This result also suggested that synthesizing BCAAs possibly confer high metabolic cost to *B. velezensis*, and relieving these metabolic burdens provide mutants with growth advantage (Fig S5A-C). It was expected that these auxotrophic mutants of *B. velezensis* SQR9 could be complemented by *P. stutzeri* XL272 during interaction. Consistently, part of the auxotrophic mutants, such as *ΔilvCH* and *ΔleuBCD* showed significantly increased cell numbers (4.3×10^5^ and 2.0×10^5^, respectively) in cocultures with *P. stutzeri* XL272 compared to monocultures of each mutant (4.4×10^4^ and 1.0×10^4^, respectively) under nutrient limited condition (MSgg medium) (Fig. 4C). Furthermore, we observed that BCAA mutants of SQR9 (*ilvD* and Δ*leuBCD*) not only benefited in cocultures, but also promoted the growth of *P. stutzeri* (Fig 4C). Nevertheless, when the mutants were co-cultivated with *P. stutzeri* in rich TSB medium, they reached similar cell numbers like the WT (Fig. S5D), likely due to excess of BCAAs in the medium. All together, these analyses indicate that *B. velezensis* SQR9 and *P. stutzeri* XL272 might exchange BCAAs when forming coculture biofilms that results in mutual benefit.

### *B. velezensis* SQR9 facilitates the growth of *P. stutzeri* XL272 by metabolic cross-feeding

To elucidate whether metabolic facilitation is responsible for syntrophic cooperation, TSB spent medium complementation assays were performed. In support of our hypothesis, the growth of *P. stutzeri* XL272 was significantly enhanced by the supplementation of *B. velezensis* SQR9 supernatant (Fig 5A), which indicated strong metabolic facilitation. However, the supernatant of *P. stutzeri* XL272 did not affect the growth of *B. velezensis* SQR9 under these conditions (Fig. 5A).

**Figure 5.**
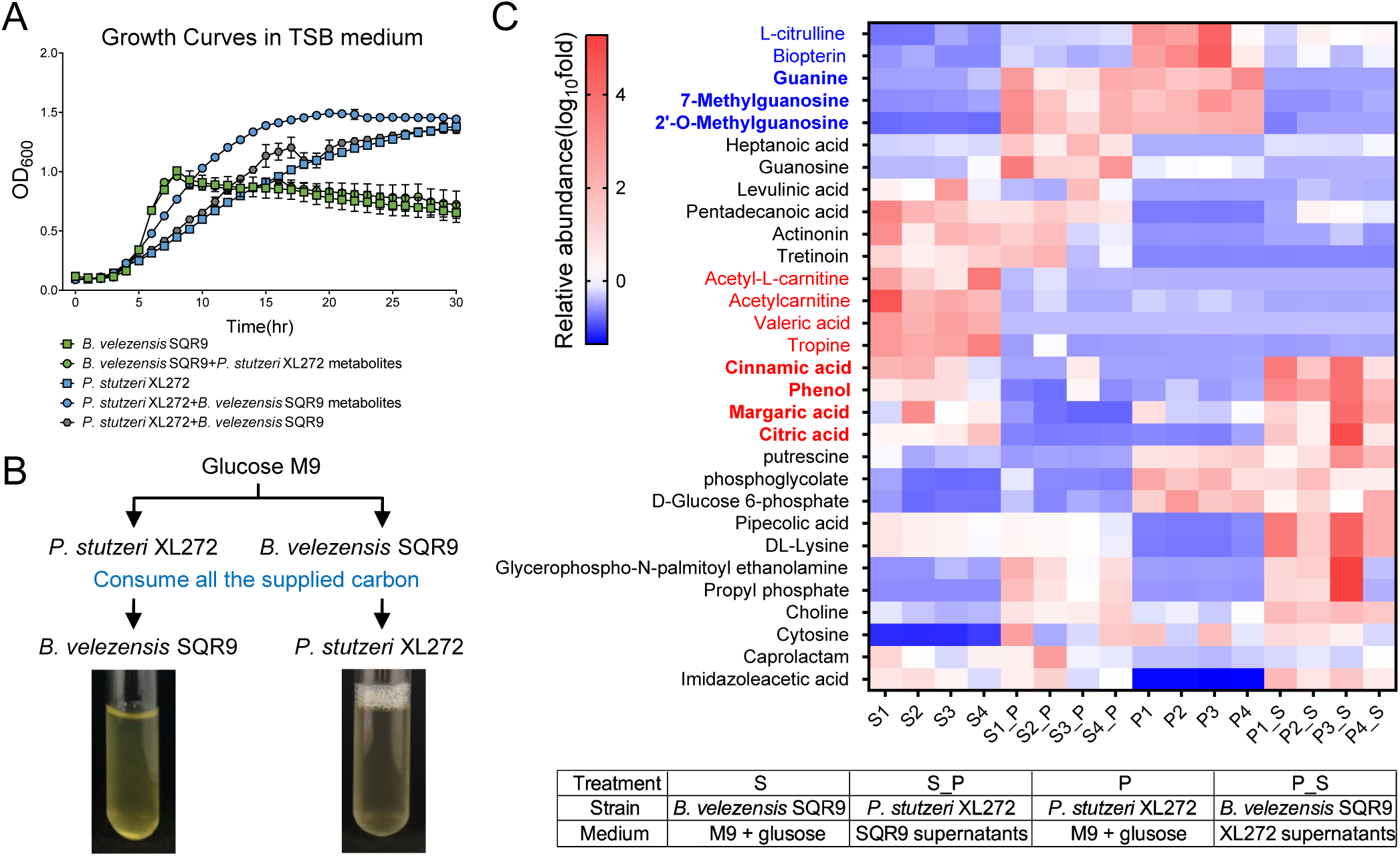
Metabolic facilitation stabilized cooperation. **(A)** Growth curves of monoculture cultivated in pure TSB media (square), monoculture grown on TSB supplemented with 10% supernatant of another species (circle) and coculture (hexagon). *B. velezensis* SQR9 TSB spent media facilitated the growth of *P. stutzeri* XL272. **(B)** Schematic representation of the experiment. Isolates were independently grown in M9 glucose medium for 6 days (*B. velezensis* SQR9) or 4 days (*P. stutzeri* XL272) till the glucose was under detection, after which cells were filtered out from the suspension. The filtrate was used as the growth media for another isolates (see methods). Both isolates were able to grow on each other isolate’s metabolic by-products. **(C)** Metabolic profiles of B. S indicated the spent medium of *B. velezensis* grown on M9 glucose, S_P indicated the spent medium of *P. stutzeri* grown on cell-free filtrate of *B. velezensis*, P indicated the spent medium of *P. stutzeri* grown on M9 glucose, P_S indicated the spent medium of *B. velezensis* grown on cell-free filtrate of *P. stutzeri*. Compounds marked blue are *P. stutzeri* metabolites that are potential preferred carbon source of *B. velezensis*. Compounds marked red are *B. velezensis* metabolites that could potentially be utilized by *P. stutzeri*.

In an alternative complementation assay, spent medium was created from M9 minimal medium with 1% glucose as sole carbon source in which monocultures of the two isolates were previously cultivated until all the supplied carbon had been consumed, thus all the carbon present in the supernatant originated from metabolites secreted by the respective bacterium. We found that both isolates were able to grow and multiply on the metabolic by-products present in each other’s, but not on their own spent medium (Fig 5B). Untargeted UPLC-MS based metabolomic approach was used to compare the metabolic profiles of cell culture grown in M9 medium and bacterial supernatant filtrate. The top 30 differential compounds were selected based on their relative abundance and displayed in a heatmap (Fig 5C). Compounds that were more abundant in glucose M9 spent medium of one species, while decreased after cultivation of the other species were assumed to be metabolized by the subsequently growing species. As a result, acetyl-L-carnitine, acetylcarnitine, valeric acid, cinnamic acid, margaric acid, citric acid, tropine and phenol were secreted by *B. velezensis* SQR9 as metabolic by-products and could potentially be utilized by *P. stutzeri* XL272 subsequently. Correspondingly, *P. stutzeri* XL272 can produce L-citrulline, biopterin, guanine, 7-methylguanosine and 2’-O-methylguanosine, which could potentially be metabolized by *B. velezensis* SQR9. It is worth noting that *B. velezensis* also produce cinnamic acid, phenol, margaric acid and citric acid (marked in bold red) after cultivated on the *P. stuteri* filtrate, and therefore these compounds could potentially be utilized by *P. stutzeri* XL272. These observations suggested that besides utilizing each other’s metabolic by-products, both species generated metabolites that were nutrient sources for the interacting partner (Fig 5C). This suggests a strong potential for metabolic cross-feeding between the two species.

The potential for metabolic cross-feeding was further supported by dissimilar preference of *B. velezensis* SQR9 and *P. stutzeri* XL272 for carbon sources that are present in root exudates (Fig. S6). Metabolic facilitation could explain the observed cooperation in TSB grown co-culture biofilms and in the rhizosphere.

### Performance of two species consortia in the rhizosphere

Since *Pseudomonas* spp. were recruited by *B. velezensis* SQR9 and the two isolates showed syntrophic exchange in biofilm formation *in vitro*, we were interested in the biological relevance of biofilm formation on plant roots. First, we monitored the colonization of these two strains on the roots of model plant, *A. thaliana*, which allows easy monitoring of bacterial colonization in the laboratory [36, 44]. *P. stutzeri* XL272 colonized the roots of *A. thaliana* at higher abundance than *B. velezensis* SQR9 using hydroponic conditions (Fig 6). Notably, these two species were able to co-colonize the *A. thaliana* roots and mixed more homogeneously than in pellicle and colony biofilms (Fig 2B, 2D &6A). We further monitored the proportion of each strain on these root-surface biofilms. In contrast to pellicle biofilm, *P. stutzeri* XL272 was the dominant species in the population colonized in the *A. thaliana* roots (Fig 6B).

**Figure 6.**
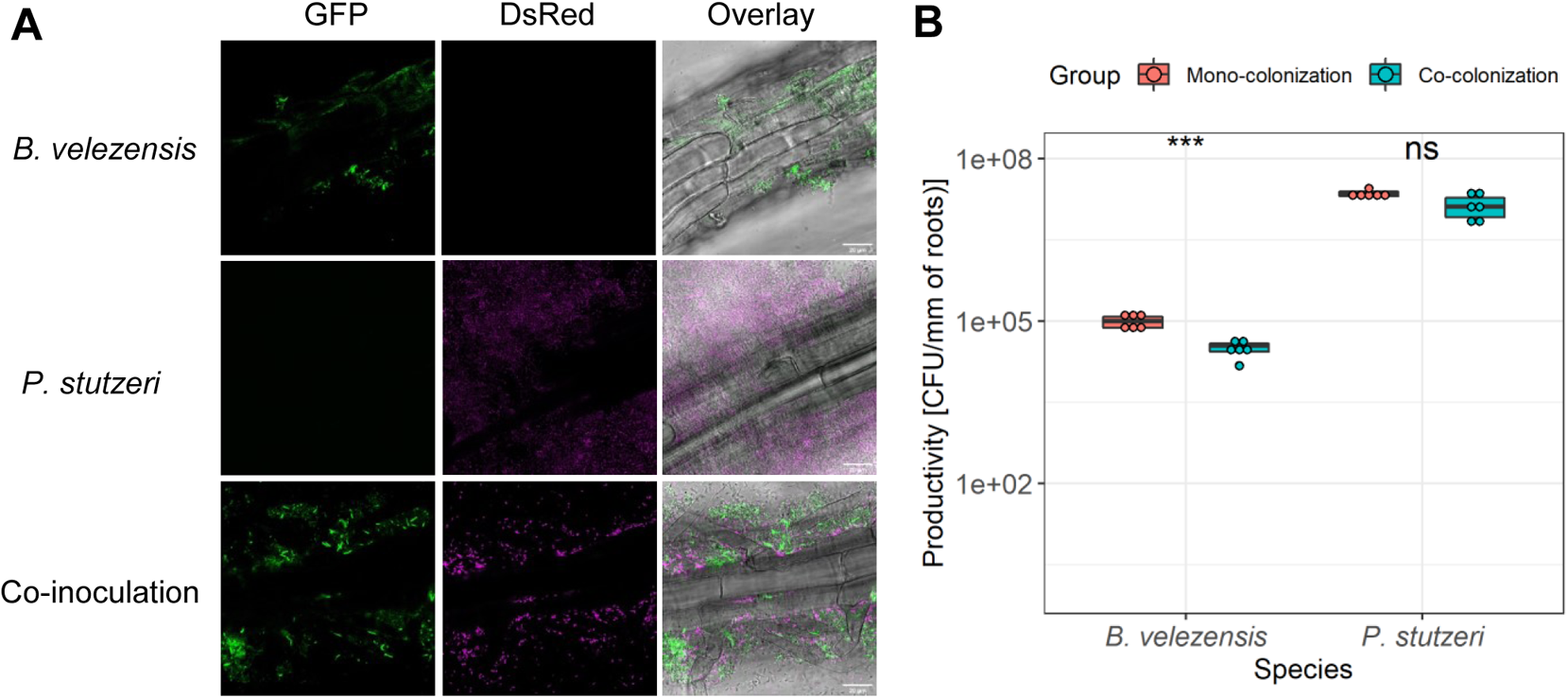
Co-colonization on plant roots. **(A)** *A. thaliana* roots were colonized by *B. velezensis* SQR9 (GFP, colored green), *P. stutzeri* XL272 (DsRed, false colored magenta) and mixture, visualized using CLSM. Scale bar represents 20 µm. **(B)** Biofilm productivities were measured as CFUs per millimeter of root (n = 6) (see Methods). The productivity of mono-colonization is compared with co-colonization. Asterisks indicate statistically significant (p < 0.01) according to unpaired student’s *t* test via R.

We further tested whether the two species Syncom performed better than the single strains in the cucumber pot assays. Before test, Indeed, results showed that both strains were able to promote the growth of cucumber plants individually and in the mixture (Fig 7). The growth promotion ability of *P. stutzeri* XL272 could be attributed to IAA, ammonia and siderophore production (Fig S7). Remarkably, the consortium had a stronger promoting effect in paddy soil, as the Syncom significantly increased the shoot height, shoot dry weight, and chlorophyll content of plants in comparison to plant inoculated with one species. In previous work, *B. velezensis* SQR9 was shown to enhance plant salt tolerance [19], the consortium was also tested under salt-treated paddy soil. Compared with non-inoculated control plants, *P. stutzeri* XL272 protects the plant against salt stress (Fig 7). The protective effect of the Syncom was higher than that of *P. stutzeri* XL272 alone in comparison to control plants. Collectively, these results showed that the genus *Pseudomonas* recruited by PGPR *B. velezensis* SQR9 in the rhizosphere promotes the growth of cucumber plants synergistically with *B. velezensis* SQR9. These results support the suggestion that the PGPR induce the assemblage of indigenous beneficial microbiome, leading to promotion of plant health and resistance to salt stress.

**Figure 7.**
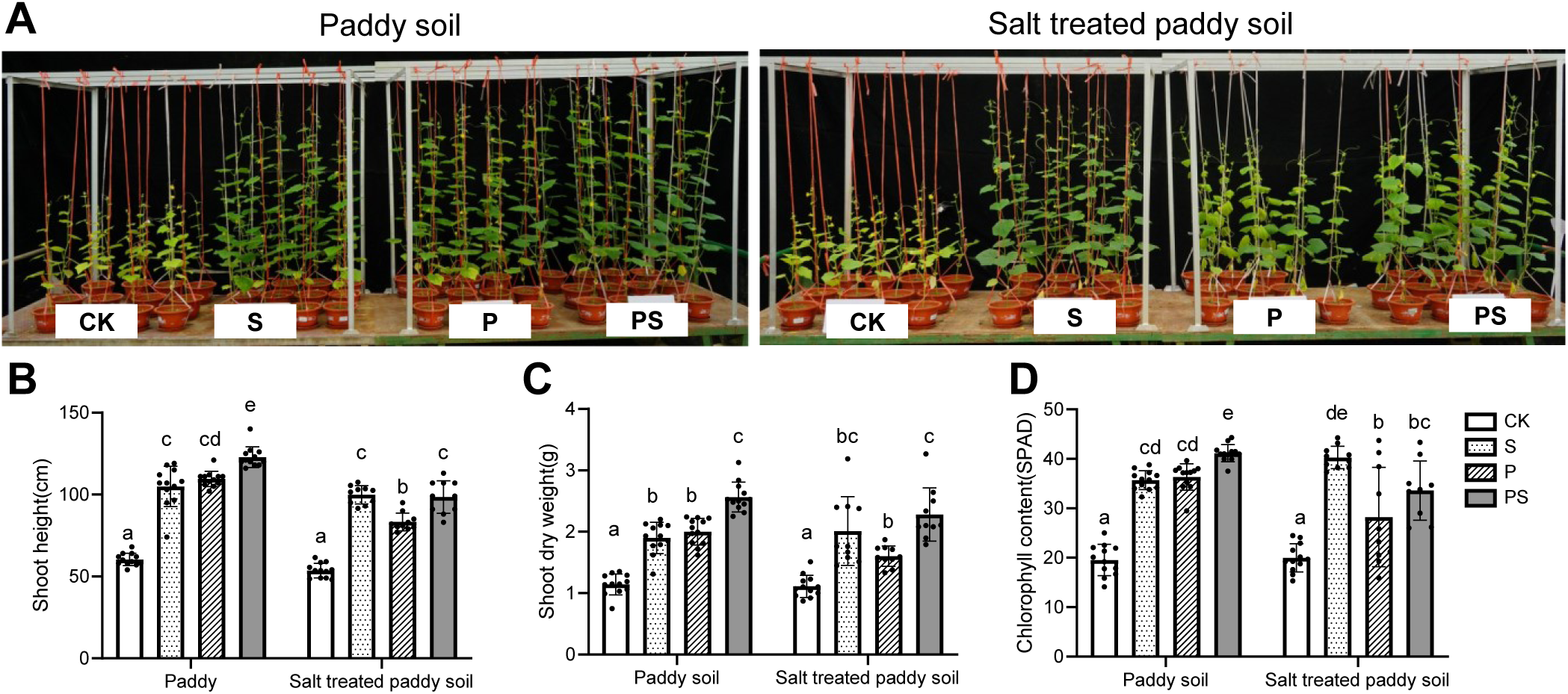
The consortium promoted plant growth and alleviated salt stress. **(A)** Photos showing 6-week-old cucumber plants grown in normal paddy soil or salt treated paddy soil inoculated with S (*B. velezensis* SQR9), P (*P. stutzeri* XL272) or PS (mixture of P and S). **(B)** Shoot height of plants. **(C)** Shoot dry weight of plants. **(D)** Leaf chlorophyll content of plants. Bars represent the mean mean ± s.d. (n = 10-12). Different letters indicate statistically significant (p < 0.05) differences according to ANOVA multiple comparison via Prism 8.

## Discussion

Just like plants employ various mechanisms to shape the composition and activity of the root-associated microbiota for their own benefits, bacterial inoculants also modulate the plant microbiota. The present study highlights the importance of microbial ecological interactions in PGPR-based biocontrol. We demonstrated that a well-established biocontrol bacterium *B. velezensis* SQR9 alters rhizosphere microbiome, by recruiting other beneficial bacteria. The recruited strains can serve as cooperative partners of SQR9 and assist in plant growth promotion.

We demonstrated that PGPR *B. velezensis* SQR9 increases the abundance of several genera with reported beneficial functions including *Pseudomonas* [45], *Chryseobacterium* [46], *Lysobacter* [47]. Stimulating beneficial bacteria in the rhizosphere seems to be a common property of *Bacillaceae*. Previously, *Bacillaceae* group species were reported to promote the colonization of rhizobia [48], promote the growth of rhizosphere bacterium *Flavobacterium johnsoniae* [49], and stimulate indigenous soil *Pseudomonas* populations that enhance plant disease suppression [50].

Here, we concentrated on the interaction between *Bacillus* and *Pseudomonas* genera as widely commercialized beneficial bacteria that dominate the rhizosphere microbiota [51, 52] and are more investigated at the molecular level compared with other genera. *P. stutzeri* XL272 was chosen as a representative to design a Syncom as it shared the same region of colonization with *B. velezensis* SQR9 in the rhizosphere and formed dual-species biofilm synergistically in TSB and REM medium. Forming biofilm synergistically is a common trait of plant beneficial consortium [41, 43]. Rhizosphere bacteria frequently reside in multispecies biofilms, the lifestyle of multispecies biofilm facilitate the emergence of community intrinsic properties [53], such as enhanced tolerance to antimicrobial agents [54, 55], horizontal gene transfer (HGT), and sharing of public goods. In this study, the dual-species community not only formed synergistic biofilm *in vitro* (Fig 2), but also created mixed species biofilms on plant roots.

We further revealed that both biofilm matrix components, EPS and TasA are important for the cooperation between *B. velezensis* and *P. stutzeri*. The role of TasA in mixed-species interaction was also reported in *B. subtilis-Pantoea agglomerans* interaction, during which both species contributed matrix components to the coculture colony biofilm structure, with *B. subtilis* producing matrix protein TasA protein and *P. agglomerans* producing exopolysaccharides [56]. In the case of *B. velezensis*-*P. stutzeri* cocultures, matrix components were important for synergism under cooperation-promoting conditions (rich medium), while under competition-promoting conditions (minimal medium), the loss of biofilm matrix conferred *B. velezensis* with a fitness advantage in the presence of *P. stutzeri*. We hypothesize that in minimal medium, *B. velezensis* might exploit the matrix components produced by *P. stutzeri*. Whether and how *P. stutzeri* complements the lacking extracellular matrix components of *B. velezensis* in the mixed species biofilms needs further investigation. Our results showed that the interaction outcome of *B. velezensis* with *P. stutzeri* is highly context dependent. The mutualism between the two species only occurred under static, nutrient-rich conditions. As static conditions allow biofilm formation, the cooperation could be facilitated by more intimate interactions between the two species, facilitating the metabolic cross-feeding, as previously demonstrated [57]. This would also be in line with aggregation-promoting matrix components being important for the cooperative interaction under this condition.

In the hydroponic root colonization assays, the environment was mixed evenly by continuous shaking of the cultures and limited plant exudates were available to bacteria, which might explain the lower colonizing cell numbers of *B. velezensis* in co-inoculation compared with that in mono-colonization. Competition is more prevalent in free-living habitats wherein the resources are scarcer, in contrast, cooperative interaction is likely to be restricted to nutritional-rich environments such as the rhizosphere. These findings are in accordance with previous studies, support the impact of spatial organization and nutrient condition on microbial interaction [58, 59].

Interaction between members of *Bacillus* and *Pseudomonas* genera was previously reported to be dependent on abiotic conditions and species involved. In particular cases these two genera utilize sophisticated competition strategies, such as type VI secretion system used by *P. chlororaphis* to attack *B. subtilis* colonies [60]. It was suggested that *B. subtilis* either produces extracellular matrix to reduce infiltration by *P. chlororaphis* or enters sporulation as self-defense strategies [60]. In another example, the presence of *B. subtilis* reduces the appearance of spontaneous mutants lacking secondary metabolite production in *P. protegens* [61]. In other circumstances, *B. licheniformis* and *P. fluorescens* interact positively in biofilm mode, enhancing plant growth and photosynthetic attributes [40], further supporting our findings.

So far, the role of *Bacilli* in recruiting beneficial bacteria was only investigated using phenotypic analysis and amplicon sequencing, with limited knowledge about interspecies interactions at the molecular level. To close this gap, we took advantage of dual RNA-seq analysis to characterize the transcriptional consequences of bacterial interactions. Transcriptome analysis revealed distinct gene expression profiles in dual-species biofilm compared with single-species biofilm. For *B. velezensis* SQR9, we noted that six pathways related to amino acids biosynthesis were downregulated in presence of *P. stutzeri* XL272, possibly reducing the metabolic costs related to these pathways. In accordance, the essential amino acids could partly be compensated by *P. stutzeri* XL272. In synthetic microbial communities, the metabolic exchange of biosynthetically costly amino acids tends to promote strong cooperative interactions [62]. This hypothesis was further confirmed using *B. velezensis* SQR9 auxotrophic mutants of BCAA biosynthesis, which could be rescued by *P. stutzeri* XL272 under nutrient-limiting conditions. Thus, the ability to synthesize BCAAs is important to survive under nutrient limited condition, while dispensable when the required nutrients are provided by co-existing species or other environmental sources.

We demonstrated that supernatants obtained using M9 minimal medium were able to support the growth of the respective other species, indicating metabolic cross-feeding, as reported previously for various mixed-species communities [10, 63–65]. Syntrophic interactions mediated by the exchange of nutrients might help to maintain the stability of the bacterial consortium. As species interactions alter the evolution of resource utilization [65, 66], it will be interesting to investigate whether the cooperation between *B. velezensis* SQR9 and *P. stutzeri* XL272 can be maintained in a longer time scale and whether the interacting partners influence the evolutionary diversification of each other.

Finally, the combined use of *Bacillus* and *Pseudomonas* inoculants provide a synergistic effect on plant health, including plant growth promotion, salt stress alleviation, as observed in our pot experiments, and increased disease suppression [50]. Inspired by their complementary colonization dynamics on plant roots [67] and positive interaction in biofilm formation, we hypothesize that fast-growing *Bacillus* could serve as a pioneer to occupy the available niche in the rhizosphere at an early stage. Through metabolism, they secrete metabolic by-products that increase the viability of successor *Pseudomonas*. In line with this hypothesis, *B. velezensis* SQR9 had a broader ability of catabolizing sugars, while *P. stutzeri* XL272 had a stronger ability to utilize organic acids (Fig S6). In agreement with the metabolome data, *B. velezensis* SQR9 convert glucose to organic acids which could be metabolized by *P. stutzeri* XL272. Whether such metabolic cooperation occur in the rhizosphere environment and contribute to the synergistic plant beneficial functions, remains to be elucidated. Basically, *P. stutzeri* XL272 was capable of IAA, ammonia and siderophore production in the lab condition (Fig S7). However, we considered that understanding the molecular processes underlying the plant-microbe interactions of *P. stutzeri* XL272 is an important step to disentangle the synergy in the rhizosphere context.

In conclusion, our findings demonstrate that a PGPR recruits indigenous beneficial bacteria and can cooperatively interact with them via cross-feeding. Synergistic biofilm formation was accompanied by enhanced plant-growth-promoting and salt stress-relieving ability. Based on our findings, the conceptual ecological biocontrol model was summarized in Figure 8. In the first step, *B. velezensis* SQR9 is attracted by root exudates and colonizes the rhizosphere [68–70]. After establishing biofilm on plant roots, it secretes metabolites that increase the abundance of indigenous plant beneficial genera (such as *Pseudomonas* spp.). By forming tightly associated biofilm, they share extracellular matrix and essential metabolites that increase their fitness in the rhizosphere. As a result, the simplified community has an increased ability to promote plant growth and alleviate salt stress. Our study proposes an ecological approach for plant health using microbial inoculants with synergistic effects, in addition to being an important step towards understanding of microbial interactions in the rhizosphere microbiome.

**Figure 8.**
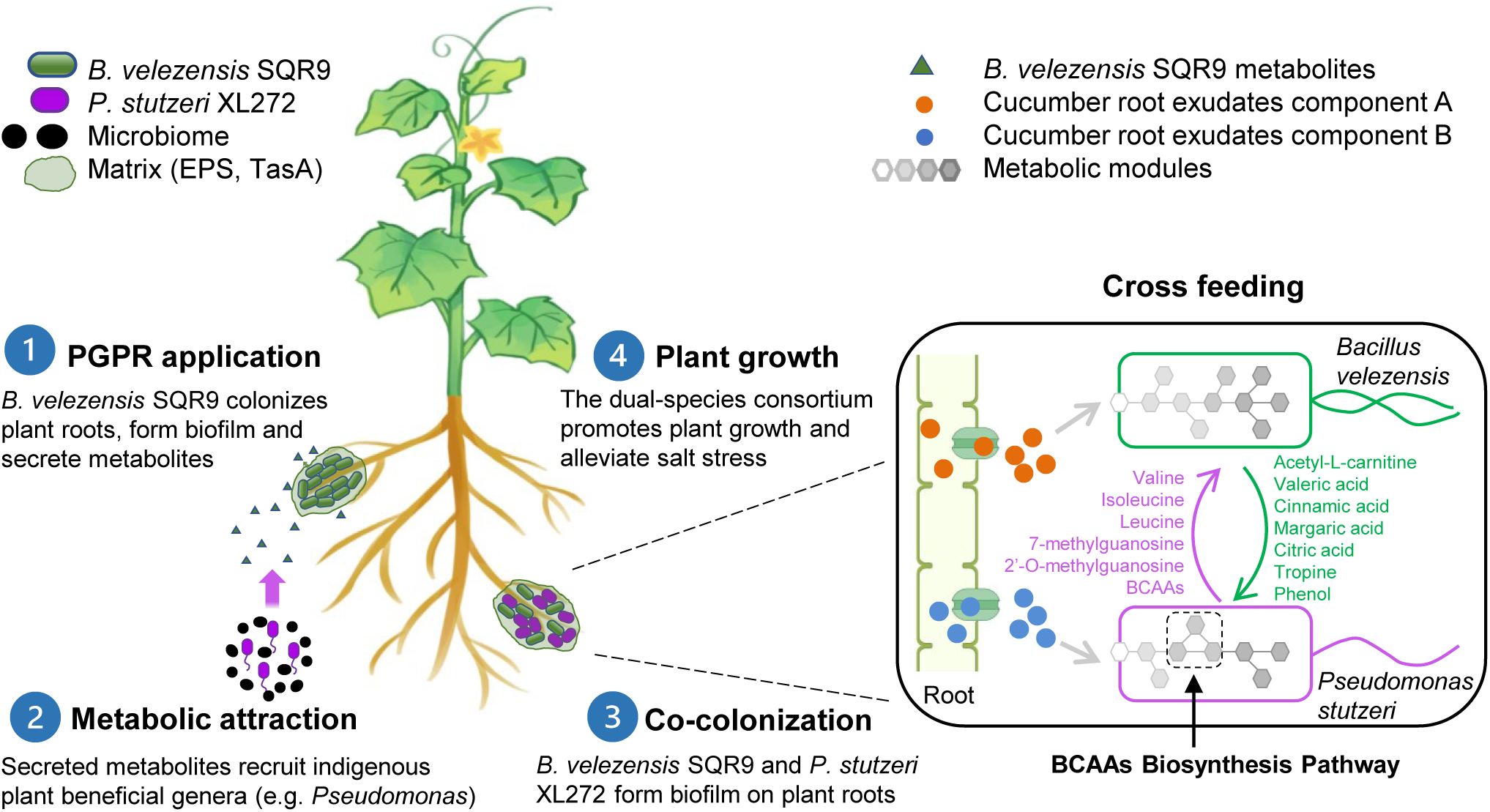
Schematic diagram illustrating the sequential events taking place in the rhizosphere after PGPR *B. velezensis* SQR9 application.

## Acknowledgements

This work was financially supported by the National Nature Science Foundation of China (31972512, 32072675 and 32072665), the Agricultural Science and Technology Innovation Program of CAAS (CAAS-ZDRW202009), the Fundamental Research Funds for the Central Universities (KYXK202009). XS was supported by a Chinese Scholarship Council fellowship. ATK, MLS and AD were supported by the Danish National Research Foundation (DNRF137) for the Center for Microbial Secondary Metabolites. Biofilm related work in the group of Á.T.K. is supported by a DTU Alliance Strategic Partnership PhD fellowship. Funding from Novo Nordisk Foundation (grant NNFOC0055625) for the infrastructure “Imaging microbial language in biocontrol (IMLiB)” is acknowledged. Author XS is very grateful to Prof. Shen and Prof. Zhang for the strong supports during the period of outbreak epidemics of COVID-19.

## Author contributions

XS, ZX, RZ, ATK designed the study, XS, JX performed the experiments. XS, VHT, MLS analyzed the data and created the figures. XS and ZX wrote the first draft of the manuscript, AD, ATK, MLS, RZ and QS revised the manuscript.

