## Supplementary material for "*Bacillus velezensis* stimulates resident rhizosphere *Pseudomonas stutzeri* for plant health through metabolic interactions"

### Supplementary figures

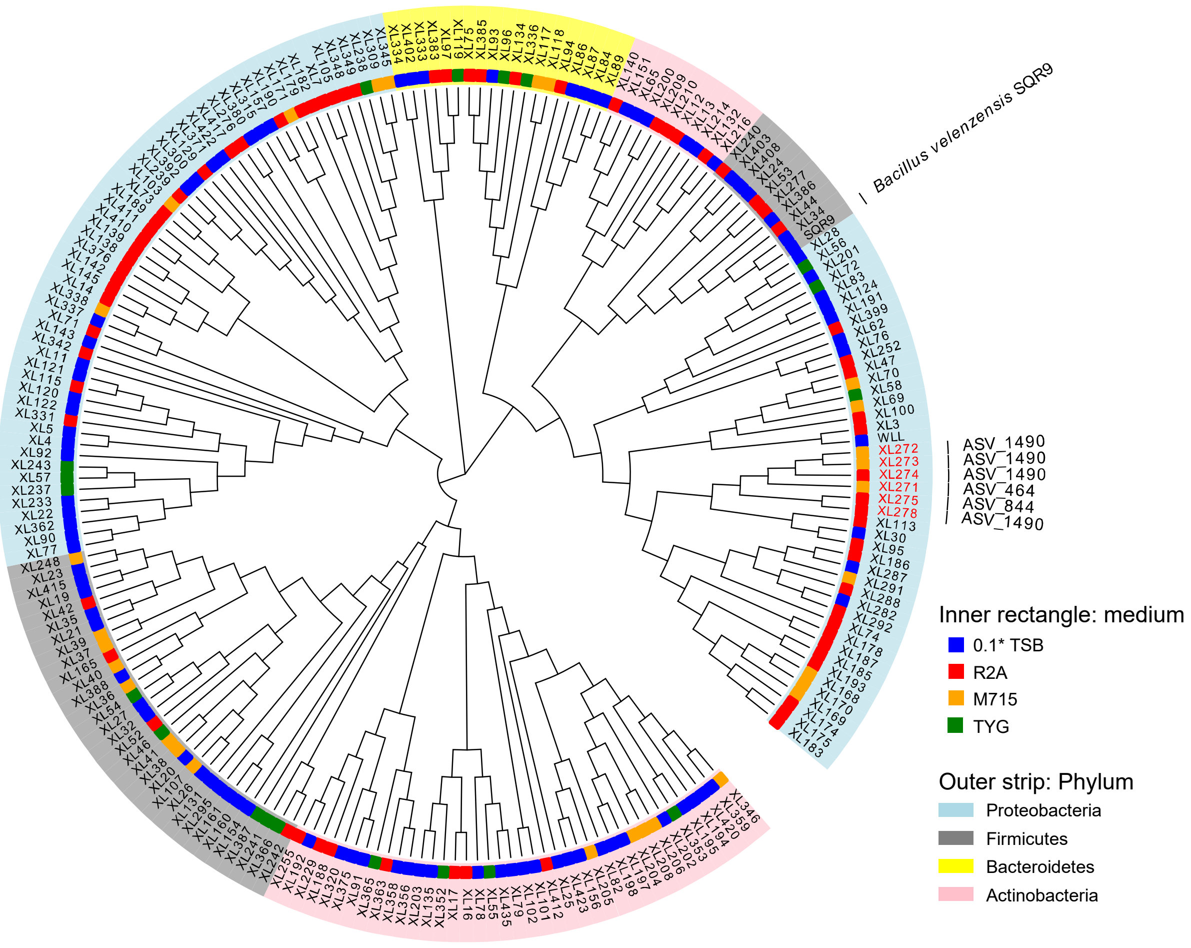


**Fig. S1 Cladogram showing phylogenetic relationships between 267 bacterial isolates obtained from cucumber rhizosphere inoculated with *B. velezensis* SQR9.** Leaf labels indicate representative sequence IDs. Labels marked red represent *Pseudomonas* isolates. Inner ring represents the medium on which isolates were originally obtained. Outer strips indicate phylum-level taxonomy of isolates. Annotation texts are the ASVs matching to the isolates with highest sequence similarity.

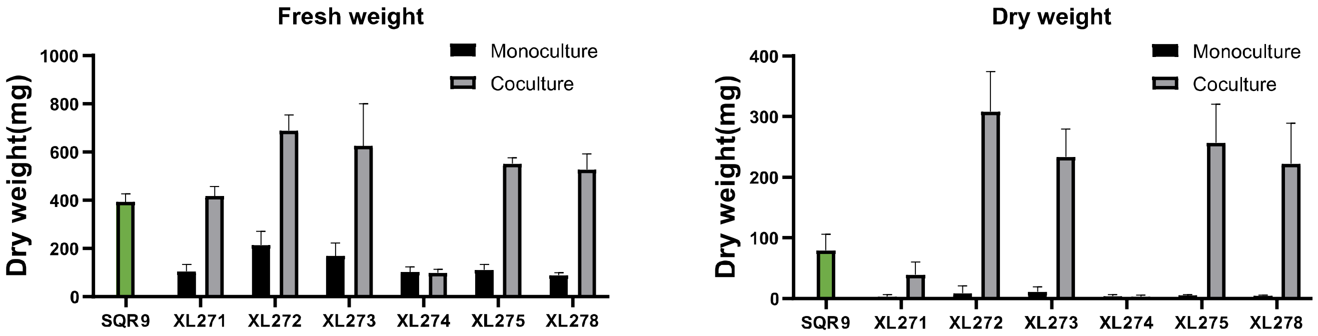


**Fig. S2 Pellicle biomass quantified by fresh weight and dry weight.** SQR9 represent *B. velezensis* SQR9, XL271-278 represent different *Pseudomonas* spp. Green bars represent monoculture of *B. velezensis* SQR9. Coculture means co-cultivated with strain SQR9. Pellicles were cultivated in TSB medium for 24h. Data presented are the mean ± s.d. (n = 4-6). Error bars represent standard deviations. Different letters indicate statistically significant ($p < 0.05$) differences according to ANOVA test via Prism 8, false discovery rate was controlled by Benjamini-Hochberg method.

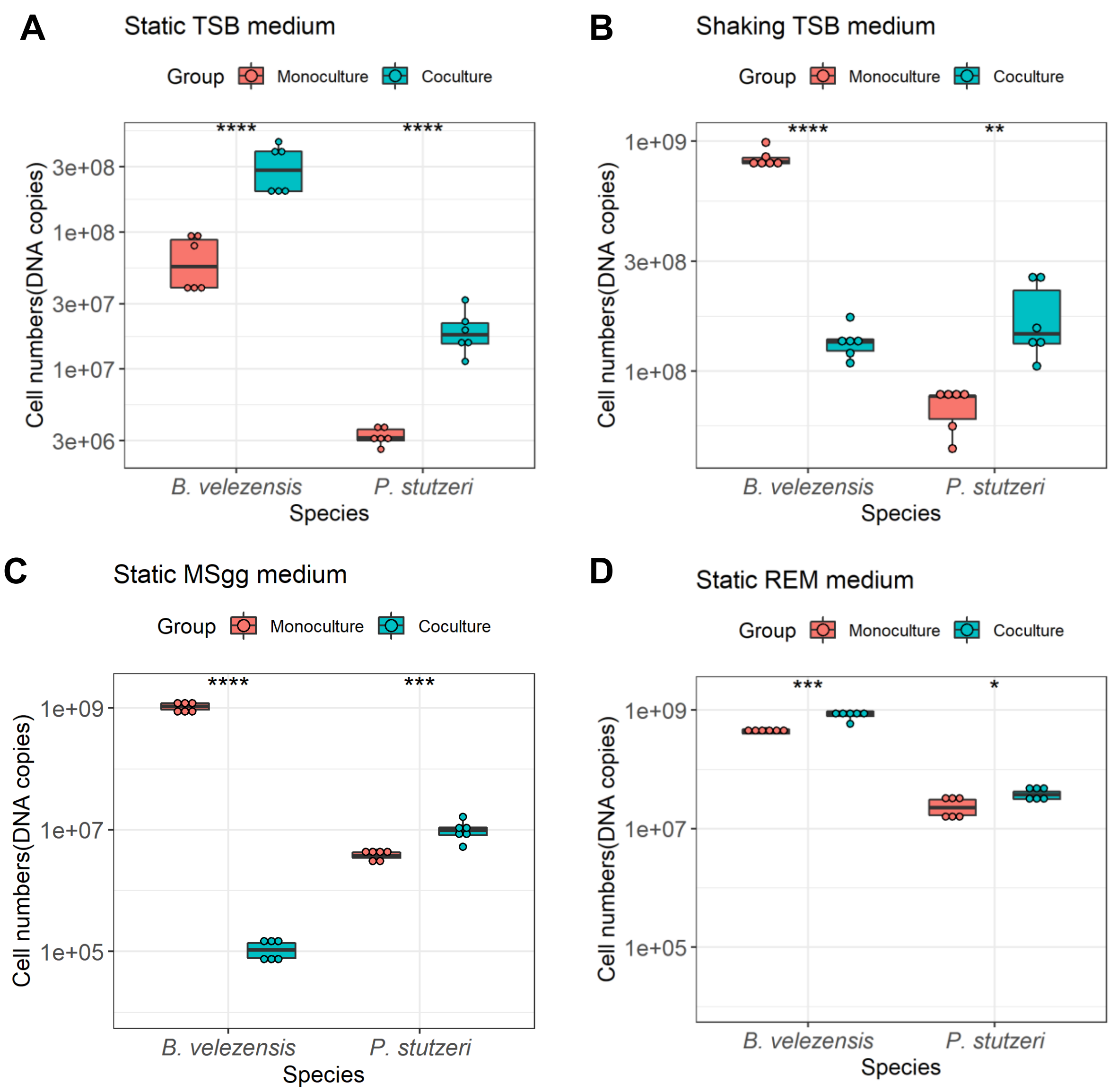


**Fig. S3 Cell numbers of *B. velezensis* SQR9 and *P. stutzeri* XL272 in monoculture and coculture.** They showed facilitation in static TSB medium, while competition in other condition. **(A)** Static TSB medium. **(B)** Shaking TSB medium. **(C)** Static MSgg medium. **(D)** Static REM medium. Asterisks indicate statistically significant ($p < 0.01$) according to unpaired student's *t* test via R.

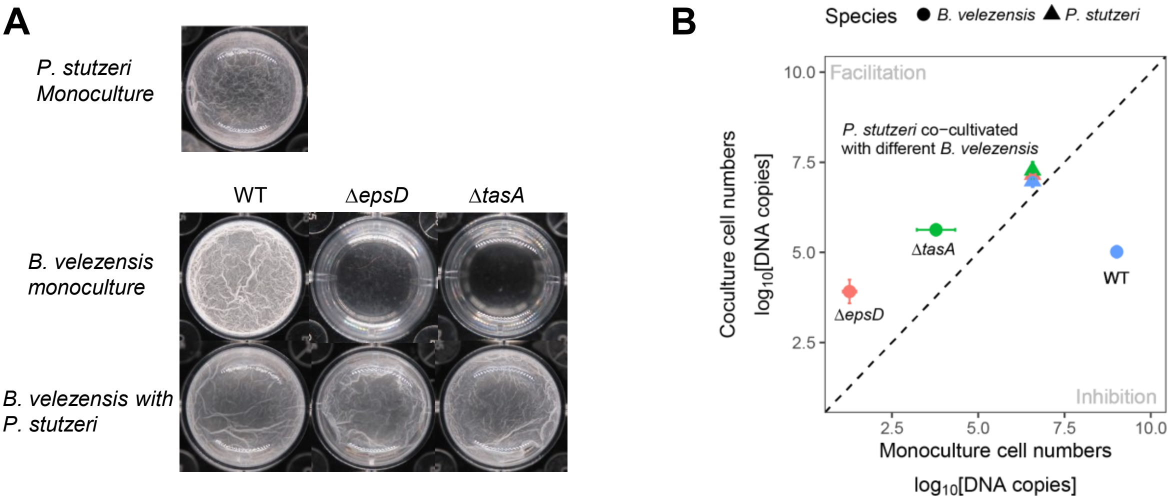


**Fig. S4**. **EPS and TasA are essential for interaction in MSgg minimal medium**. **(A)** Formation of pellicle biofilms by the mutants deficient in biosynthesis of exopolysaccharide EPS (Δ*epsD*) and TasA protein fibers (Δ*tasA*). Cells were incubated in MSgg at 30 °C for 24h before images were taken. Well diameter is 15.6 mm. **(B)** Cell numbers in dual species biofilm. Circle dots represent *B. velezensis*, triangles represent *P. stutzeri*. Colors indicate *B. velezensis* in coculture or corresponding co-cultivated *B. velezensis,* WT (blue), Δ*epsD* (pink), Δ*tasA* (green). Data presented are the mean ± s.d. (n = 6). Error bars represent standard deviations.

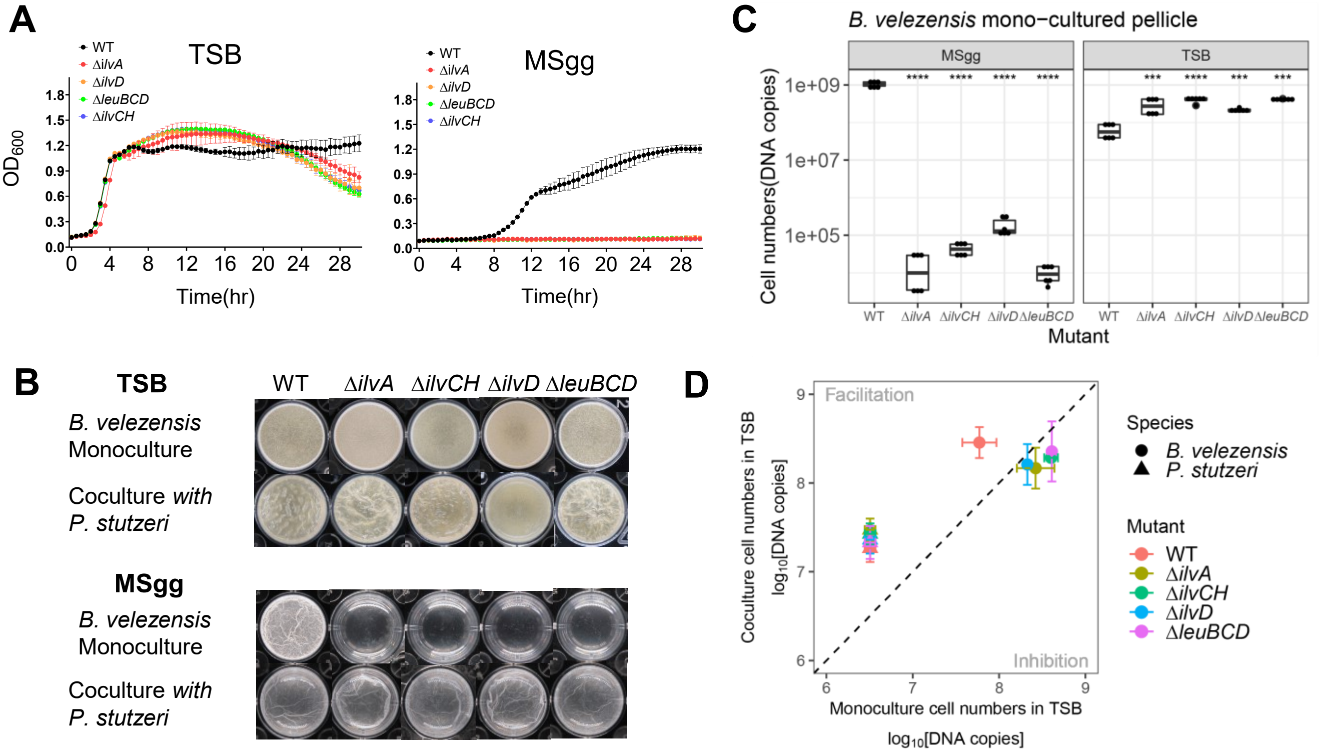


**Fig. S5 Ability to synthesize BCAAs are essential for *B. velezensis* SQR9 to survive in minimal medium but dispensable in rich medium. (A)** Growth curves of *B. velezensis* BCAA biosynthetic mutants. **(B)** Formation of pellicle biofilms by the mutants. Cells were incubated in TSB or MSgg at 30 °C for 24h before images were taken. **(C)** Cell numbers of *B. velezensis* mutants in monocultured pellicle. Asterisks indicate statistically significant ($p < 0.01$) according to unpaired student's *t* test via R. Well diameter is 15.6mm. **(D)** Cell numbers of dual species biofilm in TSB medium. Circle dots represent *B. velezensis*, triangles represent *P. stutzeri*. Colors indicate *B. velezensis* in coculture or corresponding co-cultivated *B. velezensis* WT (pink), Δ*ilvA* (brown), Δ*ilvCH* (green), Δ*ilvD* (blue), Δ*leuBCD* (magenta). *P. stutzeri* XL272 had little influence on the growth of *B. velezensis* SQR9 mutants in nutrient rich condition. Data presented are the mean ± s.d. (n = 6). Error bars represent standard deviations.

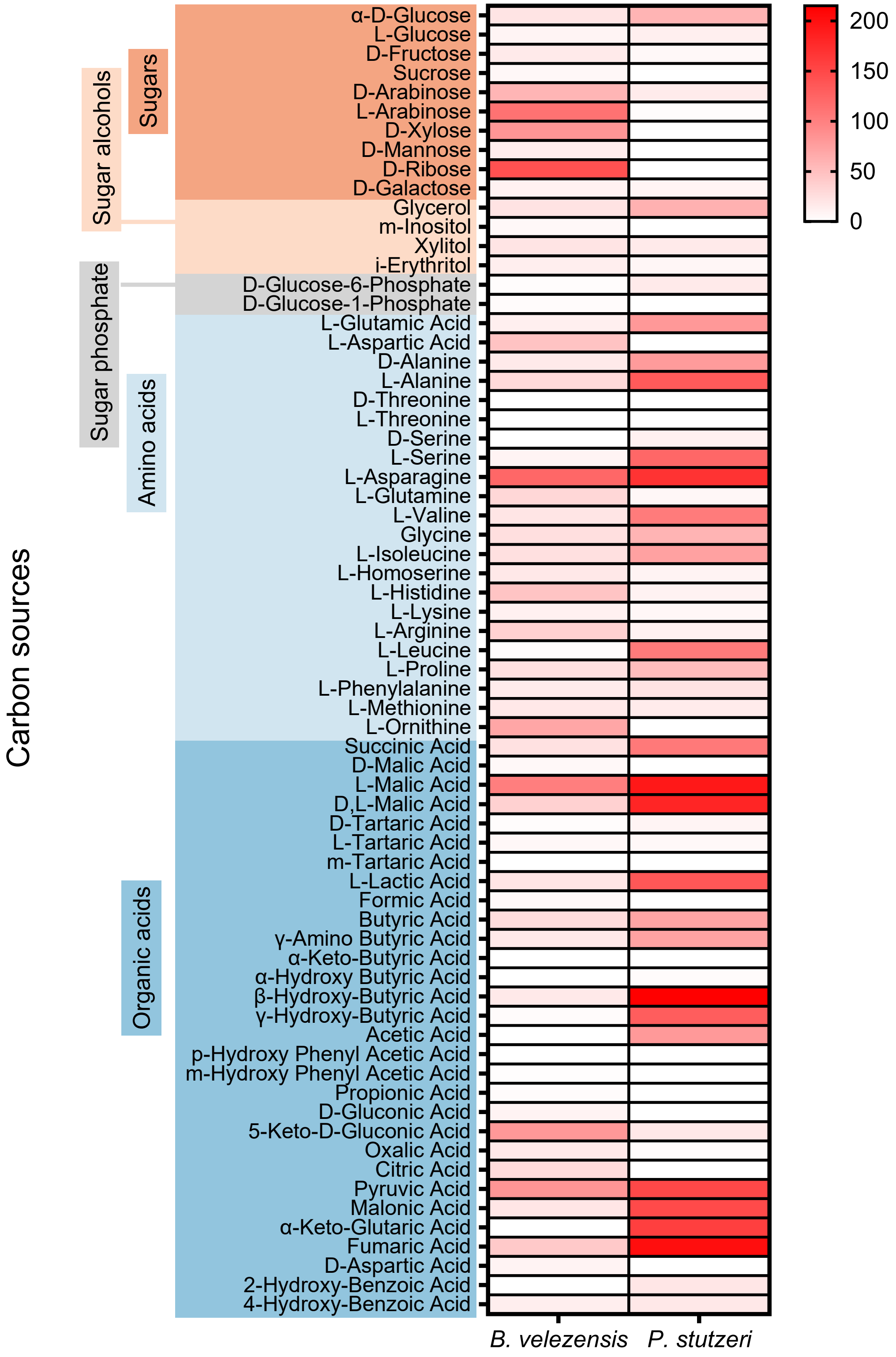


**Fig S6**. **Differential utilization of root secreted carbon sources.** Heatmap colour indicates utilizing ability determined by Phenotype Microarray(PM) technology. Root secreted carbon sources come from the literature [1].

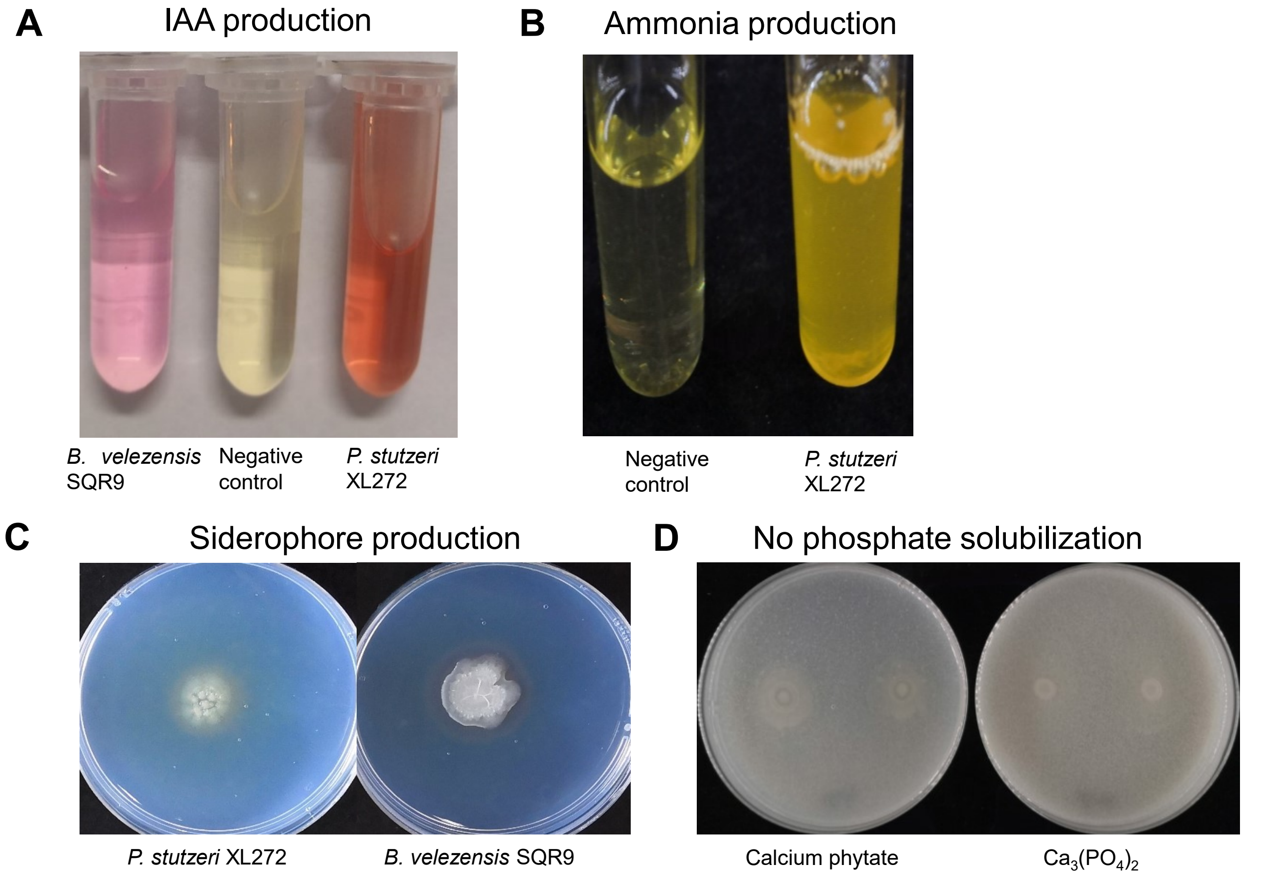


**Fig S7**. **Plant-growth promoting traits of *P. stutzeri* XL272. (A)** *P. stutzeri* XL272 is capable of IAA production at a concentration of 12.23 μg/mL. **(B)** *P. stutzeri* XL272 is tested positive for production of ammonia. **(C)** *P. stutzeri* XL272 is capable siderophore production. **(D)** *P. stutzeri* XL272 can not solubilize phosphate. Plate diameter is 9 cm.
